# Monod model is insufficient to explain biomass growth in nitrogen-limited yeast fermentation

**DOI:** 10.1101/2021.06.09.447824

**Authors:** David Henriques, Eva Balsa-Canto

## Abstract

The yeast *Saccharomyces cerevisiae* is an essential microorganism in food biotechnology; particularly, in wine and beer making. During wine fermentation, yeasts transform sugars present in the grape juice into ethanol and carbon dioxide. The process occurs in batch conditions and is, for the most part, an anaerobic process. Previous studies linked limited-nitrogen conditions with problematic fermentations, with negative consequences for the performance of the process and the quality of the final product. It is, therefore, of the highest interest to anticipate such problems through mathematical models. Here we propose a model to explain fermentations under nitrogen-limited anaerobic conditions. We separated the biomass formation into two phases: growth and carbohydrate accumulation. Growth was modelled using the well-known Monod equation while carbohydrate accumulation was modelled by an empirical function, analogous to a proportional controller activated by the limitation of available nitrogen. We also proposed to formulate the fermentation rate as a function of the total protein content when relevant data are available. The final model was used to successfully explain experiments taken from the literature, performed under normal and nitrogen-limited conditions. Our results revealed that Monod model is insufficient to explain biomass formation kinetics in nitrogen-limited fermentations of *S. cerevisiae*. The goodness-of-fit of the herewith proposed model is superior to that of previously published models, offering the means to predict, and thus control fermentations.

**Importance:** Problematic fermentations still occur in the winemaking industrial practise. Problems include sluggish rates of fermentation, which have been linked to insufficient levels of assimilable nitrogen. Data and relevant models can help anticipate poor fermentation performance. In this work, we proposed a model to predict biomass growth and fermentation rate under nitrogen-limited conditions and tested its performance with previously published experimental data. Our results show that the well-known Monod equation does not suffice to explain biomass formation.

## 1 Introduction

The yeast species *Saccharomyces cerevisiae* is the best-studied eukaryote. *S. cerevisiae* presents unique characteristics such as its fermentation capacity and its ability to withstand adverse conditions of osmolarity and low pH. These desirable properties have made of *S. cerevisiae* one of the workhorses of bio-industries including food, beverage -especially wine- and biofuel production industries (1).

In winemaking, the fermentation turns grape must into an alcoholic beverage. Wine production occurs in a closed system (i.e. in batch conditions) and is, for the most part, an anaerobic process. During fermentation, yeasts transform sugars present in the must into ethanol and carbon dioxide. A complete alcoholic fermentation is achieved when the residual fermentable sugar is less than 2 g/L. Despite improvements in fermentation control, problematic fermentations such as, stuck - with a higher than desired sugar residual- and sluggish -unusually long- fermentations still occur in real practice.

Various studies have shown that insufficient levels of assimilable nitrogen contribute to stuck or sluggish fermentations (2). A minimum of 120–140 mg/L of assimilable nitrogen is required to achieve a standard fermentation rate, while nitrogen content in grape juice may be as low as 60 mg/L (3). Therefore, it has become common practice to supplement nitrogen-deficient musts with diammonium phosphate (4). Nevertheless, both excessive or insufficient nitrogen can lead to the production of undesired metabolites affecting the organoleptic properties of wine (4, 5).

The need to predict and control wine fermentation has motivated a quest for mechanistic models of alcoholic fermentation (see the reviews by (6, 7)). Previous studies proposed various alternatives that differ on the way they explain biomass formation and the relation of cell mass (or sometimes cell numbers) to fermentation rate. Dynamic kinetic models incorporate the Monod equation to describe biomass growth (8–10). The estimation of the Monod parameters requires nitrogen and sugars uptake data throughout the fermentation. Otherwise, lack of identifiability of the parameters will limit the predictive capability of the model, and thus logistic functions may be more suitable (11, 12). Both formulations, Monod or logistic, achieve a maximum biomass value which can not be exceeded.

However, (13) showed that, for nitrogen-limited anaerobic fermentations of *S. cerevisiae*, there is a substantial increment in biomass content after the depletion of ammonia. Besides, measurements of protein and messenger RNA (mRNA) contents revealed that their concentrations stay stable while the concentration of carbohydrates increases. Similar effects have been detected in baker’s yeast exposed to nitrogen starvation but not in the carbon starved cells; see (14) for a fed-batch aerobic example, or (15) for a chemostat anaerobic example. These data would imply that, at least for ammonium deficient scenarios, the use of the Monod model would be insufficient.

Similarly, (16) showed that protein/carbohydrate fractions in the biomass of *S. cerevisiae* vary substantially throughout the fermentation of wine fermentations. To account for this dynamics, (17) and (18) included, in their flux balance analysis models, an empirical function that controls the concentration of carbohydrates in the newly formed biomass as a function of extracellular sugar.

In what regards to the modelling of fermentation rate, previous studies proposed two alternatives. The first implies that the fermentation rate is proportional to biomass or cell numbers (8–10, 12, 19). The second includes the role of hexose transporters as a separate entity that is dependent on the transport of ammonia (11, 20–23).

The advantage of the second over the first is that it enables the simulation of the effect of nitrogen additions. Still, further experimental analysis of the underlying hypothesis is required. Also, a good adjustment to the data comes at the cost of a large number of estimated parameters.

(17) and (18) proposed a third alternative. These authors assumed that cells use different hexose transporters at different fermentation stages based on previously published experimental data (24). However, this approach also requires the adjustment of the maximum transport rate for each hexose carrier. As the maximum transport rate is deemed to be a function of temperature and the initial nitrogen concentration, the modelling also involves the estimation of a substantial number of parameters.

In this work, we propose a dynamic kinetic model capable of successfully explaining nitrogen-limited fermentations. The model describes growth and fermentation rate while distinguishing protein from carbohydrate fractions of biomass. The Monod equation describes the growth, and the carbohydrate accumulation observed after the depletion of ammonia is described by an empirical function, analogous to a proportional controller.

We tested the model properties by fitting the data provided by (13) and (16) who explored fermentations under low and high nitrogen regimes. Our results show why standard Monod models often produce poor results in modelling nitrogen-limited fermentations of *S. cerevisiae*.

Fermentation rate can be a function of the biomass or, as proposed here, a function of the protein fraction since the glycolytic enzymes are an important part of the protein pool. Remarkably, even if protein content is not measured, the model is identifiable provided the uptake of hexoses and nitrogen, as well as the production of ethanol, are measured over time.

We compared the performance of the proposed model with the Monod model (25) and the model proposed by (8) who used it to explain nitrogen-limited fermentations. Our results showed that the proposed model captures the data more accurately than previously published alternatives and allowed the incorporation of macro-molecular biomass composition data (protein, mRNA and carbohydrates), which is relatively simple to obtain. The proposed mechanisms have a sound biological interpretation and were largely supported by the experimental data. In consequence, our model can be used to predict, and thus, control, *S. cerevisiae* fermentations.

## 2 Results

### 2.1 Modeling biomass formation during nitrogen-limited fermentation

The microbial growth rate was described as the sum of two different terms. The first, *μ_N_*, corresponds to the healthy cell division, mainly when assimilable nitrogen sources (YAN) are abundant. The second, *μ_C_*, corresponds to a secondary increase in biomass after the depletion of nitrogen sources- which we assumed corresponds, mostly to carbohydrate accumulation. The biomass growth equation reads:

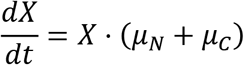

where X is the biomass (g/L).

To model growth associated with cell division we used a Monod type kinetics (25) where nitrogen sources were considered the limiting nutrient:

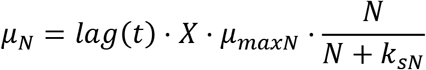

where 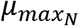 is the maximum specific growth rate, *N* is extracellular YAN (g/L), *k_sN_* is the Monod equation parameter and lag is a function of time (t) representing the lag phase. The lag phase is typically a period of adjustment in which a given microbial population adapts to a new medium before it starts growing exponentially. To model the lag we used the model proposed by (26):

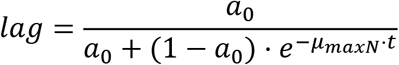

where *a*_0_ is an estimated parameter and *μ_maxN_* is the maximum growth rate associated with *μ_N_*.

The secondary increase in biomass concentration (*μ_C_*)(is activated by the of decrease the concentration of YAN:

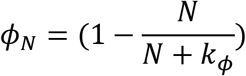

where *k_sN_* is a parameter controlling the half-maximal inactivation of *μ_N_* which is modeled with an empirical expression analogous to a proportional controller:

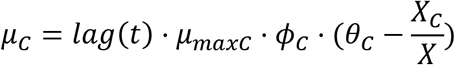

where *θ_C_* is the set-point (target value) for the ratio of carbohydrates in the biomass content (*X*_C_/*X*) and *μ_maxC_* controls the velocity of the convergence towards that reference point.

On the contrary the formation of protein and mRNA are only affected by *µ* and their dynamics are described by:

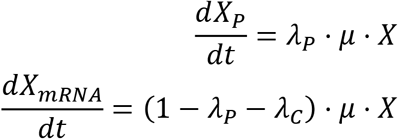

where *λ_P_* corresponds the biomass content of protein.

Both M_*M*_ and M_*P*_ assume YAN consumption is proportional to *μ_N_*:

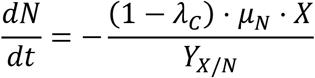

where *Y_X/N_* is the nitrogen to mRNA and protein biomass yield.

For the sake of comparison we used two additional models: the first, regarded as M_*M*_, corresponds to that for which the biomass follows the Monod description:

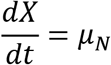

and the second, regarded as M_*C*_, corresponds to the model proposed by (8) and subsequently used by (9) to explain nitrogen-limited fermentations. In this model, the biomass follows a Monod equation [eq:Monod] depending on the active biomass and the dynamics of cell mass includes a death rate proportional to the extracellular ethanol concentration.

### 2.2 Modeling fermentation rate and production of extracellular metabolites

Regarding fermentation rate, we proposed two models with different underlying hypotheses:

- M_*P*_: assuming that fermentation rate is proportional to the total protein content (*X_P_*),
- M_*X*_: assuming that fermentation rate is proportional to the total biomass (*X*)

The uptake of glucose reads as follows:

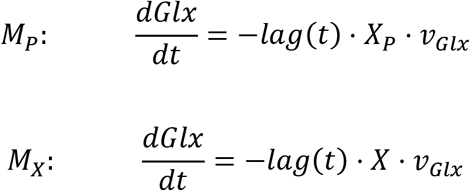

where *v_Glx_* is the expression governing glucose transport. A similar expression was also included for fructose (*F*) transport (*v_F_*):

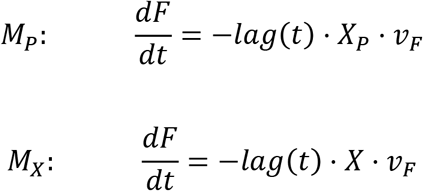

Following (27), we used Micheaelis-Menten type kinetics kinetics coupled to ethanol inhibition to model hexose transport:

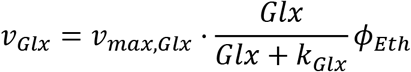

where *v_max,Glx_* is the maximum rate of glucose transport, *k_Glx_* is the Michaelis-Menten constant and *ϕ_Eth_* ethanol inhibition. Ethanol has been reported as a non-competitive inhibitor (28) of hexose transport. Here, we modeled its effect as follows (27):

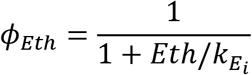

where *Eth* is the extracellular ethanol concentration and 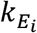 defines the strength of the inhibitory effect.

The excretion of extracellular metabolites was assumed to be proportional to the consumption of hexoses. In the case of ethanol, this is described by:

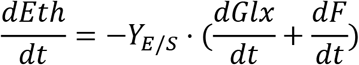

where *Y_E/S_* is the ethanol yield produced from glucose and fructose.

### 2.3 Calibration of candidate models

The Table 1 summarizes the main characteristics of the three candidate models. Additionally, a detailed description of models is given Supplementary Text S1.

**Table 1.**
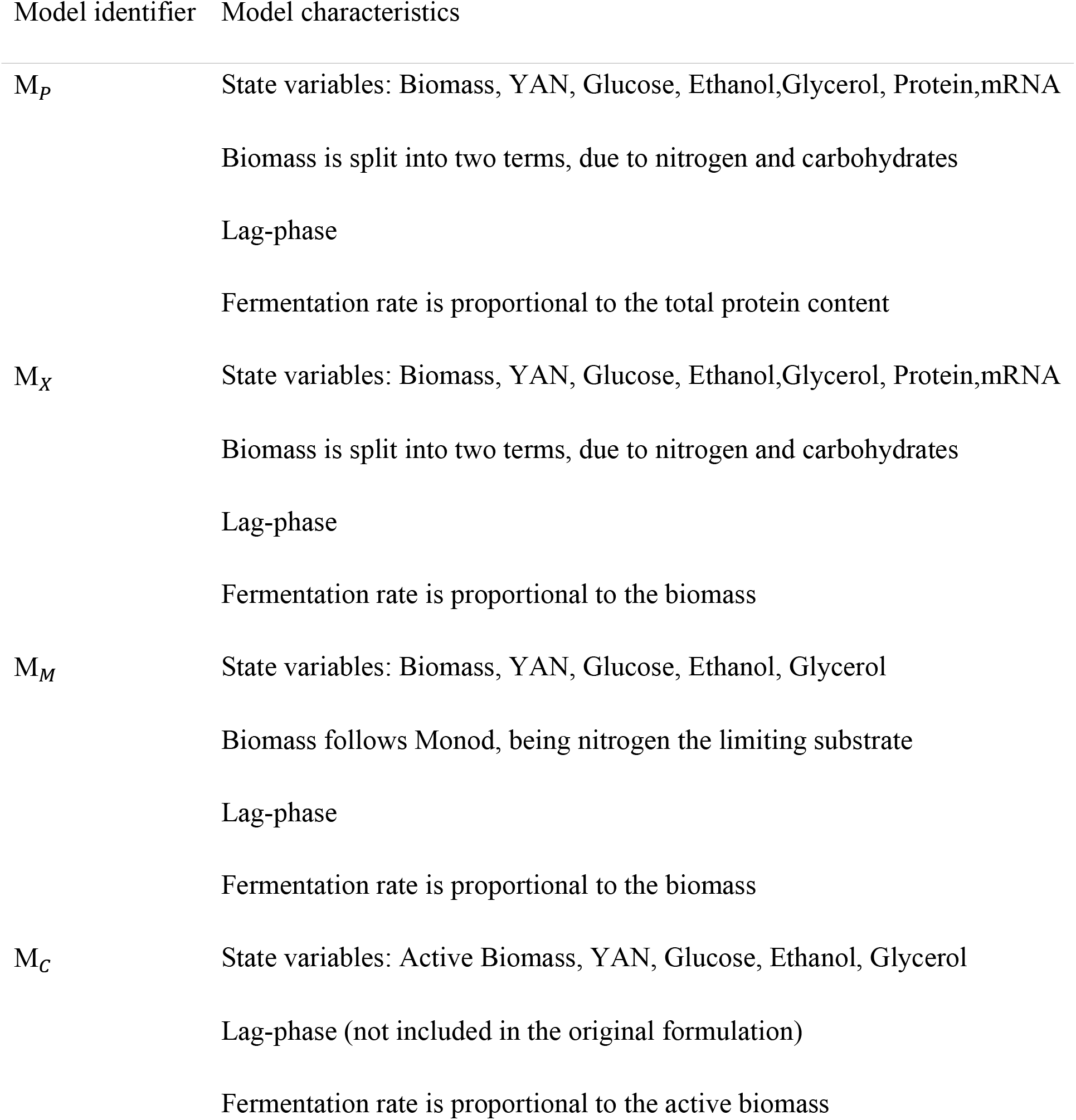
Main characteristics of candidate models. The complete mathematical formulation, including all parameters and their units are reported in the Supplementary Text S1.

The structural identifiability analysis of the models revealed that it is possible to uniquely estimate parameter values provided the dynamics of biomass, glucose uptake, nitrogen sources uptake and ethanol are measured throughout the fermentation. Remarkably, it is not necessary to measure protein for the identification of the model M_*P*_. Interested readers might find additional details in the Supplementary Text S1.

Candidate models were calibrated by data fitting using data-sets available in the literature. In particular we used data-sets taken from (13) (Exp_*S*_) and (16) (Exp_*V*_). The data-set Exp_*S*_ included one experiment with a nitrogen sufficient condition. The data-set Exp_*V*_ contained two experiments, with high (Exp_*V*_ ↑ N) and low (Exp_*V*_ ↓ N) nitrogen concentrations (50 and 300mg YAN).

We performed 10 runs of the parameter estimation problem for each case, so to guarantee convergence to the best possible solution. Since we used a metaheuristic for the optimisation of parameters, we obtained a distribution of values. We selected the best fit for ulterior analyses. Bounds for the estimated parameters values are given in supplementary text S1. Particularly, parameters related to biomass composition were chosen such that carbohydrate content does not decrease after nitrogen depletion (*i.e. λ_C_* > *θ_C_*).

The parameters recovered with the estimation procedure are given in Supplementary Text S1. Table 2 and Figure 1 present the quality of the fits in terms of normalised root mean square error (NMRSE) for each model (M_*M*_, *M_X_*, M_*P*_ and M_*C*_) and data-sets (Exp_*S*_ and Exp_*V*_). Figures 1 A-B show that all models could explain the data within a 9% normalised root mean square error. However, the performance of different models differed substantially. The model proposed in this work, M*_P_*, which explains total biomass dynamics using two terms (due to nitrogen and carbohydrates) produced NMRSE values that were 3.9 and 1.8 times lower than models M*_M_* and M_*C*_.

**Table 2.** Summary of candidate models characteristics and quality of fit scores. Observed variables regard the variables included in each model. $N_Data_ regards the amount of data used to compute the quality of fit scores. N_Pars_ regards the number of parameters to be fitted. N_RMSE_ regards the normalised mean root squared error, AIC regards the corrected Akaike criterion and Δ*AI*c the re-scaled Akaike. The table shows the model M_*P*_ results in the minimum N_RMSE_ and AIC values. Attending to the re-scaled Akaike criterion, M_X_ is the closest model to the best. However, attending to 34, models with | Δ*AIC* |>10$, have no support.

ExpS; Schulze et al. 1996
|  | MM | MX | MP | MC |
| --- | --- | --- | --- | --- |
| NData | 122 | 122 | 122 | 122 |
| NPars | 9 | 14 | 14 | 10 |
| NRMSE | 0.08 | 0.02 | 0.02 | 0.08 |
| AIC | -574.64 | -886.88 | -899.61 | -572.71 |
| $\Delta AIC$ | **-324.97** | **-12.73** | **0** | **-326.9** |

ExpV; Varela et al. 2004
|  | MM | MX | MP | MC |
| --- | --- | --- | --- | --- |
| NData | 99 | 99 | 99 | 99 |
| NPars | 10 | 16 | 16 | 11 |
| NRMSE | 0.05 | 0.04 | 0.02 | 0.05 |
| AIC | -564.74 | -597.38 | -672.47 | -564.52 |
| $\Delta AIC$ | **-107.73** | **-75.09** | **0** | **-107.95** |

**Figure 1.**
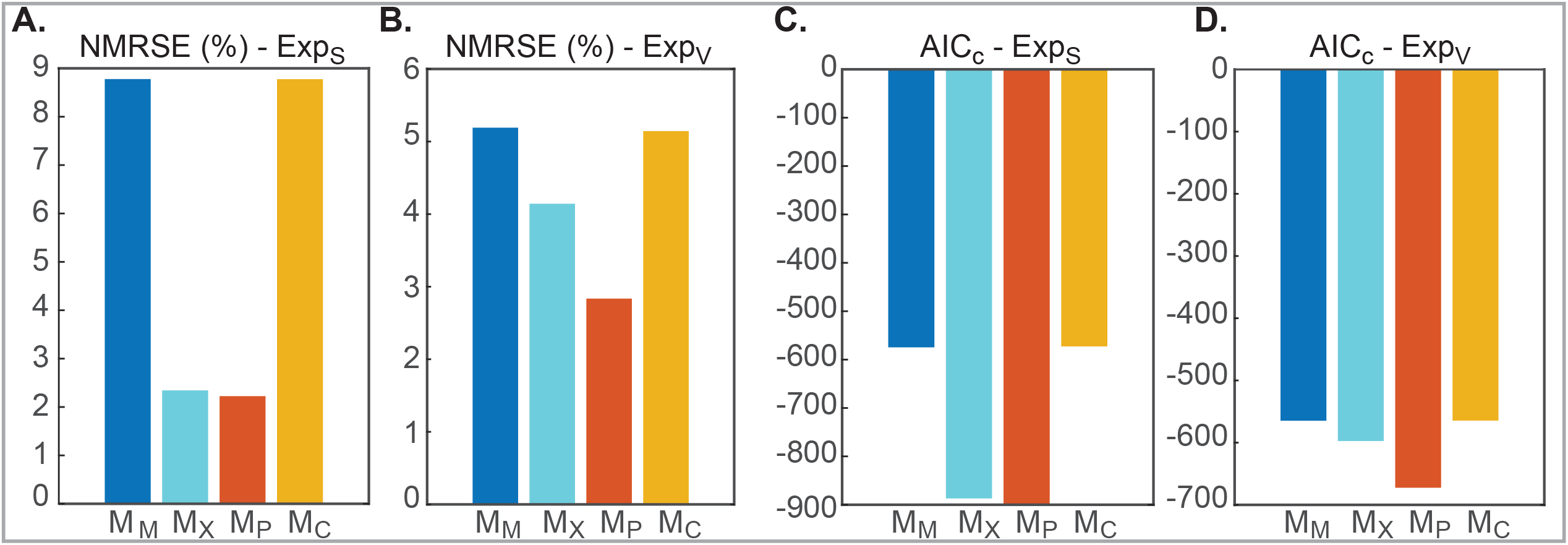
Comparison of candidate models. A-B Show the normalised root mean square error (NRMSE in %) o C-D present the corrected Akaike Information Criterion (*AIC_c_*). Values correspond to all models and the two experimental data sets Exp_*S*_ and Exp_*V*_. For the case of *AIC_c_* only YAN, glucose, fructose, ethanol and glycerol residuals were compared, since those are the variables present in all models. Results show how the proposed model M*_P_* is superior in terms of both scores for both experimental data sets, differences are particularly important in the fit to Exp_*V*_ in which high and low nitrogen fermentations are fitted simultaneously.

Since the proposed models (M*_P_* and M*_X_*) have more parameters than M_*C*_ and M*_M_*, we computed the corrected Akaike Information Criterion (AIC) to rule out any possible over-fitting. We computed AIC scores excluding the residuals corresponding to protein, carbohydrates and mRNA – as these variables are not included in M*_M_* and M_*C*_– and considering the total number of parameters of each model (including *λ*_C_,*λ_P_*, *k_sC_* and *μ_maxC_*). Table 2 and Figures 1 C-D present the results. The model M*_P_* presented substantial differences (< −100) with respect to M*_M_* and M_*C*_. Remarkably these differences are enough to guarantee that the extra parameters, used to characterise biomass composition are not inducing over-fitting and that this model is indeed the best to explain the data.

### 2.4 Biomass formation

Our results suggest that Monod model is insufficient to explain biomass formation kinetics in nitrogen-limited fermentations of *S. cerevisisae*. Figure 2.A presents the NRMSE obtained for biomass in all data-sets and models. Results illustrate the proposed model M*_P_* represents better the data in all cases. The largest differences between models were found in the Exp*_S_* data-set. Remarkably, the normalised root mean square error obtained for the proposed model (NMSRE (M*_P_*):1.91%) was approximately 9.7 times lower than the one obtained with the models M*_M_* and M_*C*_ (NMSRE (M*_M_*):18.60% and (M_*C*_):18.56%, respectively). For the data-set Exp*_V_*, differences were lower; still, the model (M*_P_*) proposed in this work was always superior in the quality of fit to the biomass data.

**Figure 2.**
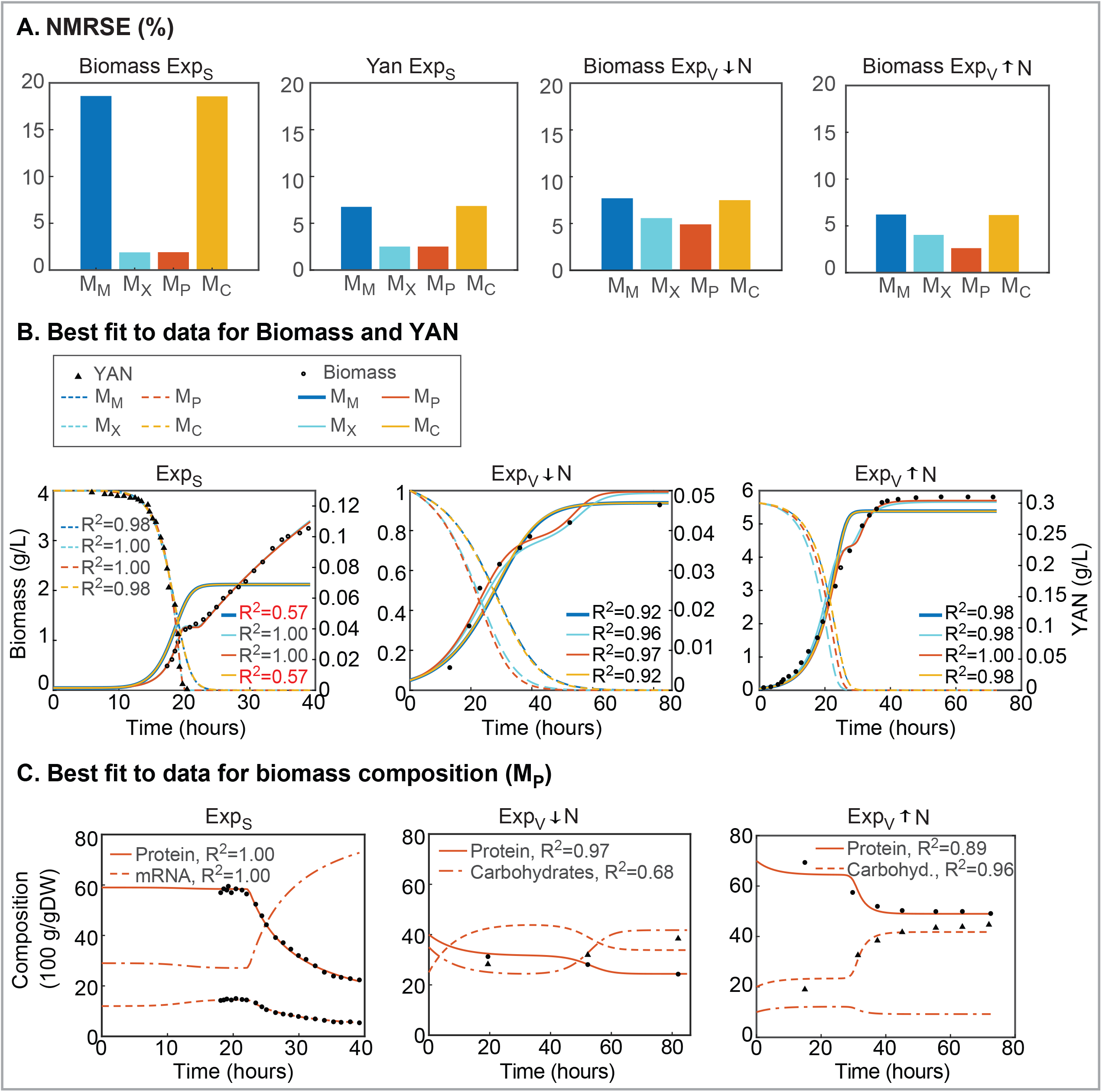
Comparison of candidate models in the fitting of biomass data. Figure A presents the normalised root mean square error of dry-weight biomass and YAN in all candidate models and experiments. Figure B presents the time-course trajectories and data for YAN and dry-weight biomass in all experiments. Figure C presents the time-course trajectories as obtained by model M*_P_* of the percentage of protein (100 ⋅ *X_P_*/*X*), mRNA (100 ⋅ *X_mRNA_*/*X*) and carbohydrates (100 ⋅ *X*_C_/*X*) present in the biomass. Data-set Exp*_V_* is comprised by experiments Exp*_V_* ↑ *N* and Exp*_V_* ↓ *N* which were fitted in combination and share the same parameters. In contrast Exp*_S_* corresponds to a single experiment fitted in isolation. Model M*_P_*, proposed here, results in better fits for dry-weight biomass in terms of NRMSE and *AIC_c_*. The model can recover the increase in biomass observed in several experiments after YAN depletion.

Figure 2.B presents the fit to the time-series data for all three considered experiments. In all three cases, the proposed model could recover the increase of biomass produced after YAN depletion. The improvement was more notorious in the Exp*_S_* data-set. Figure shows *R*^2^ values for the proposed model are over 0.97 in all cases while M*_M_* and M_*C*_ resulted in *R*^2^ of around 0.57 for the Exp*_S_*.

Figure 2.C shows how after nitrogen depletion, at around 20 hours (see also Figure 3.A, carbohydrates accumulate and would explain the role of *μ*_C_ in models M_*P* and M*_X_*. Remarkably those models are able to recover the secondary growth observed in the biomass dynamics while M*_M_* and M_*C*_ converge to a constant biomass value.

**Figure 3.**
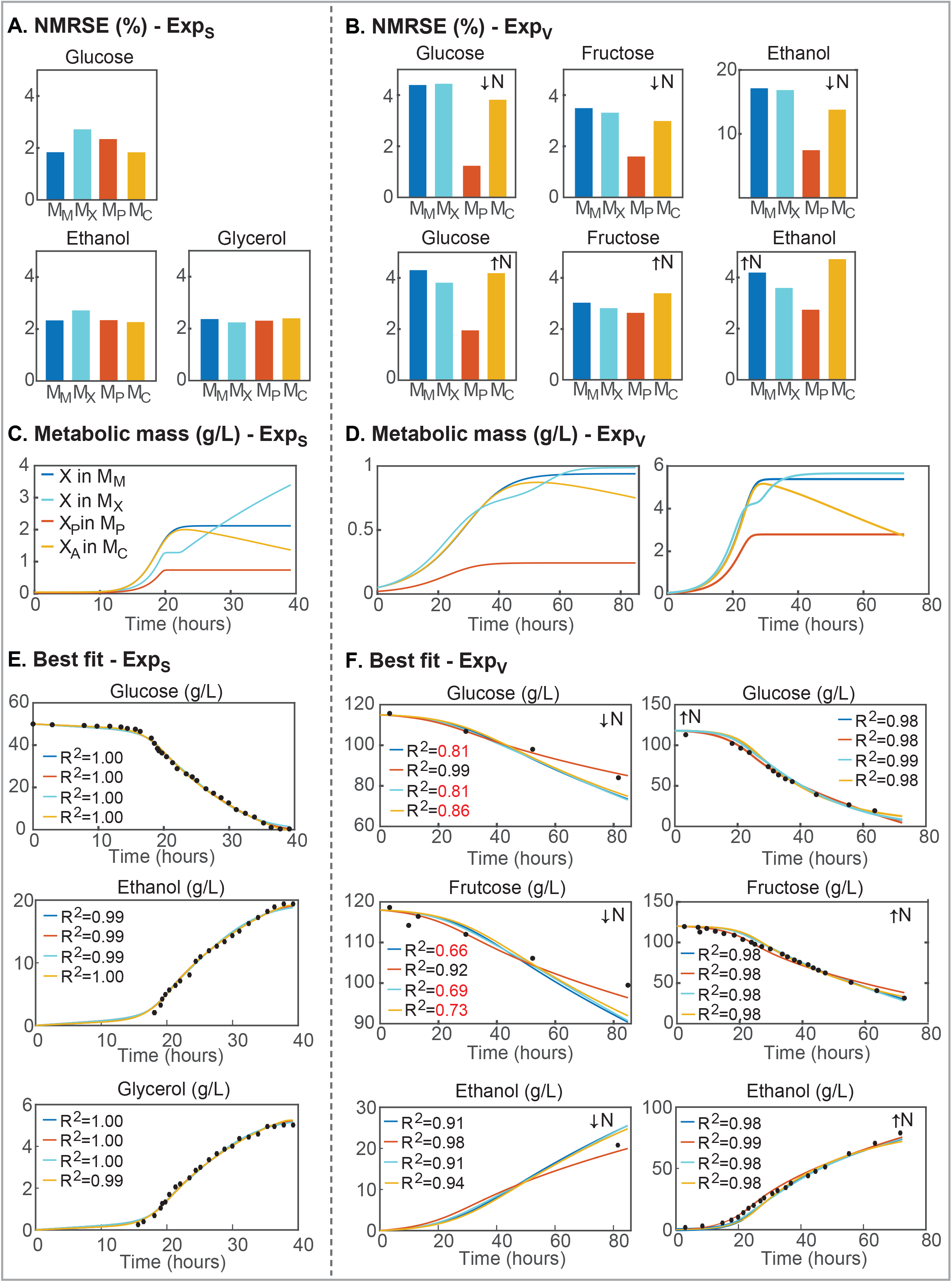
Comparison of candidate models in the fitting of fermentation data. Figures A and B present the NRMSE for the measured external compounds for the different experiments and models; Figures C and D present the time-course trajectories for the metabolically active cell mass (*X*, *X_A_* or *X_P_*); Figures E and F present the time-course trajectories of the measured variables in all experiments. *R*^2^ values are reported for each observable; those values below 0.9 are marked in red. Note that all models are quite successful in fitting the experimental data for Exp*_S_* while higher differences are observed for experiments Exp*_V_*. Only, M*_P_* resulted in *R*^2^ values above 0.9 for all states and conditions.

### 2.5 Fermentation rate

All models, were able to recover the overall dynamics of hexose consumption and ethanol formation. Figure 3.A and B show the NRMSE scores for the measured variables (glucose, fructose, ethanol or glycerol) in the different experiments. Because both M*_X_* and M*_P_* performed well in describing biomass and share the same number of parameters, a comparison is straight-forward. Remarkably the performance of the model M*_P_*, which uses the protein instead of the biomass to explain the fermentation rate, was always superior to that of M*_x_*.

Figures 3.C and D show the trajectories for metabolically active cell mass for each candidate model. Models M*_M_* and M*_X_* do not distinguish between active and total biomass, thus curves shown in 3.C and D coincide with those in Figures 2.B). Model M_*C*_ includes a decay term that may be observed in 3.C and D. In model M*_P_* active biomass is proportional to *X_P_*. As shown in 3.C and D for M*_X_* the metabolically active cell increases after YAN depletion while in M*_M_* and M*_P_* it stays stable; and in M_*C*_ there is an observable decay.

Figures 3.E and F show the trajectories for all candidate models and the corresponding experimental data for measured external compounds. Results show that all models perform reasonably well describing the dynamics of glucose, ethanol and glycerol with *R*^2^ > 0.8. However, all models but M*_P_* experiment difficulties in fitting the data Exp*_V_* with low nitrogen. In this condition M*_X_* (and also M_*C*_, M*_M_*) presented glucose consumption and ethanol productions rates faster than those observed in the experimental data; an effect not observed in M*_P_*, indicating that protein content is a helpful biomarker while modeling fermentation rate across multiple experimental conditions.

## 3 Discussion

In this study, we proposed two candidate models to explain nitrogen-limited fermentations of *S. cerevisiae* in industrially controlled conditions. We proposed two modifications to the standard modelling approaches: 1) the biomass growth accounts for protein and carbohydrates, and 2) fermentation rate is proportional to the total protein content. We reconciled candidate models to previously published data.

Our results revealed that Monod model is insufficient to explain biomass formation kinetics in nitrogen-limited fermentations of *S. cerevisisae*. This is so because, during exponential growth, the biomass composition seems to remain unaltered with mRNA and protein comprising an important percentage of the biomass. However, this is not the case during secondary growth phase where *μ_N_* fades and *μ*_C_ increases, as shown in (13) data.

Several recent studies on the modelling of alcoholic fermentation of *S. cerevisisae* relied on logistic growth (12, 20), Monod or the use of kinetic constraints coupled to a stoichiometric model (which in practice holds similar results to Monod).

Logistic growth is well suited to explain cell numbers in nitrogen-limited fermentations. However, as pointed by (25), biomass and cell numbers are not necessarily equivalent. The author argued that the average size of cells varies considerably from one phase to another in the growth cycle, thus cell concentration and bacterial density, are not equivalent. However growth rate is the same independently of being estimated in terms of one or other variable. Also in the view of (25), much confusion has been created because this important distinction has been frequently overlooked. Indeed, the author highlights that in most of the experimental problems of bacterial chemistry, metabolism, and nutrition, the significant variable is bacterial density while cell concentration is essential only in problems where division is actually concerned, or where knowledge of the elementary composition of the populations is important.

Thus, in light of our results, we argue the logistic curve model might also not adequately describe the delay during the transition from exponential growth (*μ_N_* ≫ *μ_C_*) due to carbohydrate accumulation (*μ*_C_ ≫ *μ_N_*) and the different dynamics between *μ_N_* and *μ*_C_ observed in cell mass data.

From a practical point of view, nitrogen-limited Monod models seem to explain biomass formation until the nitrogen source is depleted but struggle to explain later stages. The formulation of M*_P_* implies that carbohydrates accumulation is solely responsible for the biomass increment after nitrogen depletion. In the case of Exp*_S_* and Exp*_V_* ↑ *N*, the *R*^2^ scores for YAN, dry-weight biomass, protein and mRNA are in line with the notion that biomass increment in secondary growth phase could be explained mostly by carbohydrate accumulation.

The models proposed by (8, 9) distinguished active from inactive cells. The model M_*C*_ (adapted from (8)) was able to explain fermentation rate when data was fitted to a single experiment. However, when the same model was applied to a data-set comprised by two different nitrogen conditions (Exp*_V_* ↓ *N* and Exp*_V_* ↑ *N*) the same set of parameters had difficulties explaining hexose consumption of the low nitrogen experiment.

(20) developed a fermentation model for oenological conditions that accounted for different temperatures and initial nitrogen concentrations. These authors estimated the number of hexose transporters present in the yeast cells as a function of time, temperature and nitrogen. The model represents glucose consumption data extremely well. However, it should be noted that it comes at the cost of a heavily parameterized and nonlinear hexose transport function that also includes inhibition by substrate and ethanol. Nevertheless, the notion that hexose transporters vary with nitrogen content along the fermentation is likely to find some overlap with our simpler model for glucose transport.

(17) included the effect of ethanol inhibition on sugar transport and the existence of different hexose transporters with different maximum transport rates and substrate affinities. From a qualitative point of view, this model corresponded to a significant improvement over past models. Noticeably, the hexose transport functions considered the role of the initial nitrogen concentration and temperature. Additionally, (16) showed that the protein content of biomass is heavily dependent on the initial nitrogen concentration which supports the notion that hexose transport is likely to be better described by total protein content rather than biomass *per se*. Our model would support this hypothesis. Results obtained by M*_P_* model are superior to those obtained with the M*_X_* model, at least, for those cases in which initial nitrogen is restricted.

Here we proposed an alternative modelling strategy which using a minimal number of parameters, that can be estimated from easily measured data, provides a considerable improvement in the description of nitrogen-limited fermentations as compared to previously published models. It should be noted that the model can be easily applied to controlled fermentations -with a single starter- provided the grape must is well characterized, i.e. YAN and carbon sources are measured beforehand and their values are followed throughout fermentation, and biomass growth and desired products are monitored over time.

## 4 Materials and Methods

### 4.1 Data

We considered data-sets from five different experiments as published in the literature. The first experiment (Exp1) data set was obtained from (13). The authors conducted a nitrogen-limited (165 mg NH_3_ and 50g of glucose) batch fermentation while measuring biomass, protein content of biomass along with several extracellular metabolites (glycerol, ethanol and ammonia). We also considered four data-sets from (9). The authors performed various experiments at different conditions. In this work we denote Exp2 as that performed at 11°C, with high-sugar and low-nitrogen; Exp3, the one performed at 15°C, with normal sugar and low-nitrogen; Exp4, the one performed at 30°C with normal sugar and normal nitrogen; and Exp5, the one performed at 35°C with high-sugar and low-nitrogen.

### 4.2 Model building

The proposed model is composed of six ordinary differential equations (as presented in the section Results). Solutions for the system are of type:

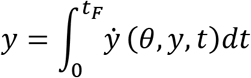

where *y* is the solution of the ODE system and corresponds to the dynamics of the relevant state variables - biomass, carbohydrates, protein, glucose, YAN, ethanol, glycerol- between the beginning and the end of the experiment 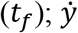 represents the states time derivative.

### 4.3 Structural identifiability analysis

The proposed model depends on fourteen unknown parameters **θ** - namely *a*_0_, *μ_maxN_*, *k_sN_*, *μ_maxC_*, *K*_sc_, *θ_C_*, *λ*_C_, *λ_P_*, *Y_X/N_*, *v_max,Glx_*, *k_Glx_*, 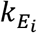, *k_Eth_*, *k_Gly_* - to be estimated from the experimental data. The proposed model introduces the description of the total protein content. However for some experimental set-ups protein was not measured. Therefore we performed a structural identifiability analysis ((29, 30)) to assess if all parameters of the model can be estimated even if protein measurements are not available.

We performed the analysis using GenSSI2 toolbox (31). The toolbox uses symbolic manipulation to implement the so-called generating series of the model and to compute identifiability *tableaus*.

### 4.4 Parameter estimation

The aim of parameter estimation is to compute the unknown parameters - growth related constants and kinetic parameters - that minimize the weighted least squares function which provides a measure of the distance between data and model predictions (32). In the absence of error estimates for the experimental data a fixed value for each state, the maximum experimental data, was assumed in order to normalize the least squares residuals

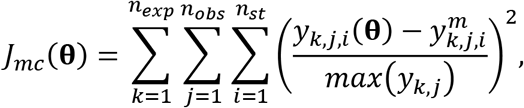

where *n_exp_*, *n_obs_* and *n_st_* are, respectively, the number of experiments, observables, and sampling times. 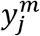 represents each of the measured quantities and *y_j_*(**θ**) corresponds to model predicted values.

In our particular case, we used five different experiments: the first, taken from (13), and second to fifth, taken from (9). All experiments provided time series data of the biomass (*X*), glucose uptake (*Glx*), amonia consumption (*N*) and ethanol production *Eth*.

Parameters were estimated by solving a nonlinear optimization problem to find the unknown parameter values (**θ**) to minimize *J_mc_*(**θ**), subject to the system dynamics - the model- and parameter bounds (33).

To avoid the risk of premature convergence by the optimization routine while searching for the optimal parameter set, the parameter estimation procedure was repeated 10 times for each candidate model starting from different initial guesses.

### 4.5 Model selection

Models were compared in terms of the Akaike’s information criterion (AIC). AIC compares multiple competing models (or working hypotheses) in those cases where no single model stands out as being the best. AIC is calculated using the number of data (*n_d_*), the number fitted parameters (*n_θ_*) and the sum of squares of the weighted residuals (WRSS). For the purpose of model comparison, we computed the AIC as follows:

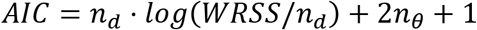

The models were ranked by *AIC*, with the best approximating model being the one with the lowest *AIC* value. The minimum AIC value (*AIC_min_*) was used to re-scale the Akaike’s information criterion. The re-scaled value *ΔAIC* = |*AIC_i_* − *AIC_min_*| was used to assess the relative merit of each model. Models such as *Δ* ≤ 2 have substantial support, models for which 4 ≤ *Δ* ≤ 7 have considerably less support and models with *Δ* > 10 have no support (34).

### 4.6 Numerical methods and tools

The parameter estimation and model selection were implemented in the AMIGO2 toolbox (35). The system of ODEs was compiled to speed up calculations and solved using the CVODES solver (36), a variable-step, variable-order Adams-Bashforth-Moulton method. The parameter estimation problem was solved using a hybrid meta-heuristic, the enhanced scatter search method (eSS, (37)).

## 5 Supplemental Material

All scripts and data required to reproduce results and figures can be accessed in https://sites.google.com/site/amigo2toolbox/examples.

## 6 Acknowledgments

We thank the three anonymous reviewers for their comments on the original version of the manuscript. This project has received funding from MCIU/AEI/FEDER, UE (grant reference RTI2018-093744-B-C33) and GAIN Xunta de Galicia (Contrato Programa and grant reference IN607B 2020-03).

